# Western Amazon was a center of Neotropical fish dispersal, as evidenced by the continental-wide time-stratified biogeographic analysis of the hyper-diverse *Hypostomus* catfish

**DOI:** 10.1101/2021.06.03.446980

**Authors:** Luiz Jardim de Queiroz, Xavier Meyer, Yamila P. Cardoso, Ilham A. Bahechar, Raphaël Covain, Thiago E. Parente, Gislene Torrente-Vilara, Paulo A. Buckup, Juan I. Montoya-Burgos

**Author notes:** Correspondence: Juan I. Montoya-Burgos. Department of Genetics and Evolution, Faculty of Sciences, University of Geneva. Quai Ernest Ansermet 30, CH-1205 Geneva, Switzerland.

## Abstract

The Amazon is probably the most diverse realm on Earth, and is considered to be the primary source of diversity and a center of dispersal for Neotropical terrestrial organisms. Yet, the assumption that the Amazon basin is a primordial place of fish species origination and dispersal into other drainages still need to be tested. We addressed this issue by inferring a time-stratified biogeographic history and reconstructing the ancestral habitat preference of *Hypostomus*, a continentally widespread and species-rich Neotropical genus. We found that *Hypostomus* emerged in the Western Amazon (∼14.7 Ma), when the Western Amazon River was flowing northwards and disconnected from the Eastern Amazon. We show that dispersal events in the first half of *Hypostomus* evolution occurred from the Western Amazon into adjacent basins, initiating its Neotropical radiation. The ancestral preferred habitat consisted in small rivers with running waters, a predominant habitat in river headwaters. Because of strong niche conservatism in the early evolution of *Hypostomus*, we suggest that most of the out-of-Western-Amazon dispersal occurred via headwater captures. The radiation of *Hypostomus* was further promoted by major reconfigurations of river basins, which opened dispersal opportunities into new drainages. Diversification in habitat preference coincided with colonization of basins already occupied by congenerics, indicative of niche shifts triggered by inter-specific competition and species coexistence. By analyzing the evolutionary history of *Hypostomus*, we show that Western Amazon was the main center of fish dispersal in the Neotropical Region from Middle Miocene to the present, supporting the cradle hypothesis of fish origination and dispersal.

## Introduction

The Neotropical Region, a large biogeographic realm that covers an area from Southern Mexico to Argentina (Cox, 2001; Leroy et al., 2019), harbors a vast variety of tropical biomes and habitats, and an unparalleled biotic diversity (Hughes, Pennington, & Antonelli, 2013). The processes that have given rise to this diversity are complex, and the many hypotheses of tempo and mode of diversification are still controversial (reviewed in Antonelli, Ariza, et al., 2018). Yet, there is a consensus on the importance of the dynamic geology and climatic variations in shaping the diversity observed nowadays (Antonelli, Ariza, et al., 2018; Antonelli & Sanmartín, 2011). Therefore, taking these processes into account is central for a better comprehension of the origin of the Neotropical biodiversity and the geographical patterns of distribution.

One of the most striking features along the history of the Neotropical Region was the uplift of the Andes, which was the result of a gradual thickening of the Earth’s crust due to the interaction between the Nazca and South American plates (Jordan et al., 1983; Pilger, 1984). The Andes is a continuous mountain range along the western edge of South America, extending over more than 7,000 km in length. This mountain range has been uplifting slowly over the last 30 million years, and the modern configuration and elevation were achieved about 14–10 million years ago (Evenstar, Stuart, Hartley, & Tattitch, 2015; C. Hoorn et al., 2010; Pilger, 1984). One of the most emblematic landscape reconfigurations driven by the Andes uplift was the modification of the course of the Amazon River (C. Hoorn, Guerrero, Sarmiento, & Lorente, 1995). Until around 10 Ma, the drainages of the current Amazon and Orinoco rivers formed together the proto-Amazon-Orinoco system (PAO), which was the most important basin that existed from the early Oligocene to the middle-Miocene in the Neotropical Region. PAO flowed northwards into the Caribbean Sea through the outlet of the contemporary Lake Maracaibo (C. Hoorn et al., 2010; Jaramillo et al., 2017). During the Miocene, marine transgressions changed the landscape of part of the Neotropical Region, including PAO, where a vast saline swamp-like environment was formed, the Pebasian Sea (Wesselingh, Guerrero, Rasanen, Romero Pittman, & Vonhof, 2006). With the progressive uplift of the Northern Andes, the outlet of PAO was gradually closed, blocking large masses of water in the inland. Concomitantly, the rise of a topographic elevation in Northwestern South America, the Vaupés Arch, caused the division of the POA into the western Amazonian basin and the Orinoco basin (C. Hoorn et al., 2010). Then, with the breach of the Purus Arch, which was the drainage divide separating current western and eastern Amazonian drainages, the Amazon River eventually reached the Atlantic ocean and achieved its modern configuration (James S. Albert et al., 2018; C. Hoorn et al., 1995; Carina Hoorn et al., 2017).

Following these drastic modifications of the Neotropical landscape during the Miocene, Amazonian plant and animal lineages experienced an explosive diversification (C. Hoorn et al., 2010). There is strong evidence that the Amazon region is likely the most important source of diversity of terrestrial lineages worldwide (e.g. flowering plants, ferns, birds, mammals and reptiles), playing a pivotal role in shaping the biodiversity across the American continent (Antonelli, Zizka, et al., 2018; Musher, Ferreira, Auerbach, McKay, & Cracraft, 2019). The reasons why Amazonia is a primary source of terrestrial biodiversity are probably multiple, and some interdependent features have been evoked such as the high level of environment heterogeneity spanning over a very large area, a dynamic landscape where diversification has been kept at relatively high levels and constant over time, with low extinction rates, and a high connectivity with neighboring biomes, facilitating dispersal of terrestrial lineages (Musher et al., 2019).

As for fish, the Amazon Basin is the biogeographic core of the Neotropical system, and by far the most species-rich river network (J. S. Albert & Reis, 2011; Böhlke et al., 1978), with more than 7’000 species estimated (Lévêque, Oberdorff, Paugy, Stiassny, & Tedesco, 2008; Reis, Kullander, & Ferraris, 2003). The Amazon Basin has also been suggested to be a cradle of fish diversity, an area where species have originated and immigrated into other basins (J. S. Albert & Reis, 2011; James S. Albert et al., 2018; Fontenelle, Marques, Kolmann, & Lovejoy, 2021; Oberdorff et al., 2019), but this hypothesis still needs further testing. Whether the many and profound landscape changes that occurred in the Amazon Basin were instrumental in fostering the origination, dispersal and diversification of modern Neotropical fish lineages is an assumption that needs to be verified too.

The increasing knowledge about the historical modifications of Neotropical basins offers a timely opportunity to infer freshwater fish ancestral distributions and dispersal routes integrating the changes in watershed configurations through time, but this has been explicitly explored in surprisingly few biogeographic reconstructions of Neotropical fish (e.g. Fontenelle et al., 2021; Wendt, Silva, Malabarba, & Carvalho, 2019). To take advantage of the accumulated knowledge of basins connectivity through time, it is important that reconstructions use the finest possible biogeographic partitions of the hydrological systems, but this implies extensive knowledge about species distribution as well as the inclusion of a comprehensive taxonomic sampling. When based on a limited number of biogeographic areas, as in most of the biogeographical studies of Neotropical fish (Mariguela, Roxo, Foresti, & Oliveira, 2016; e.g. Roxo et al., 2014; Silva et al., 2016), ancestral distribution ranges tend to be unrealistically large, some covering half of South America, thus losing credibility and biological relevance.

In the present work, we investigated of the role played by the various Neotropical river basins, integrating their historical changes in connectivity, as centers of origin and diversification of new fish lineages and as dispersal platforms from which neighboring basins were colonized. To this aim, we reconstructed the biogeographic history of one of the most species-rich genera in the Neotropical Region, the armored catfish genus *Hypostomus* (Loricariidae). *Hypostomus* are bottom-dwelling fishes with an herbivorous, detritivorous or xylophagous diet, and are generally territorial and poor swimmers. By considering a large fraction of the species diversity of this genus and by including representatives from most South American drainages thanks to the comprehensive phylogeny of *Hypostomus* and closely related genera we recently published [26], we assembled a dataset with one of the finest geographical resolution to date, at the Neotropical level. To work efficiently at the finest spatial resolution, we developed a new tool for the program RevBayes (Höhna et al., 2016) which is an interactive environment for Bayesian statistical analyses with a phylogenetic focus. The new tool we present here allows the user to disregard unrealistic distribution ranges when performing biogeographic analyses, accelerating the analyses and allowing the inclusion of more and finer geographical areas.

Taking advantage of our comprehensive dataset of the exceptionally diverse and widespread genus *Hypostomus*, and using it a model Neotropical fish group, we tested whether (i) the genus *Hypostomus* originated in the Amazon Basin, and (ii) the Amazon Basin was the most important source of diversity for this lineage in the Neotropical Region. Furthermore, we investigated how habitat preference evolved along the phylogeny of *Hypostomus* and how it might have influenced the dispersal pattern of this genus across the Neotropical river basins.

## Material and Methods

### Phylogenetic reconstruction and tree calibration

To reconstruct the phylogenetic tree of *Hypostomus*, we used the dataset published by (Jardim de Queiroz et al., 2020), which includes six loci: the mitochondrial (i) Cytochrome c oxidase subunit I protein-coding gene (*COI*) and (ii) *D-loop*, and the nuclear (iii) Gene encoding Teneurin transmembrane protein 3 (*Hodz3*), (iv) a *Hypostomus* anonymous marker, possible gene *ZBTB10* intron 3 (HAM-*ZBTB10-3*), (v) the *Recombination activating gene 1* (*RAG1)* and (vi) the Fish *Reticulon 4* (*RTN4*). This dataset contained 206 *Hypostomus* individuals organized in 108 putative species and 49 species belonging to closely related genera (outgroups). We completed this dataset with three additional species: two samples of *Hypostomus* sp. ‘Araguaia’ (Tocantins-Araguaia Basin, Brazil), one of *Hypostomus* sp. ‘Aqua75’ (Paraguay River, Brazil) and one of *Hypostomus annectens* (Rio Dagua, a Pacific coastal river of Colombia). The primers listed in (Jardim de Queiroz et al., 2020) were used to amplify the desired six loci for the new samples. Genbank accession numbers of the additional samples can be found in Supporting file 3 (Table S3).

The tree inference and calibration was performed using BEAST 1.8.1 (Drummond & Rambaut, 2007), while the input file (an Extensible Markup Language, XML, file) was built in BEAUTi 1 (Drummond, Suchard, Xie, & Rambaut, 2012a). We partitioned the data by gene (six partitions), and substitution models and clock models were unlinked among the partitions. A Yule process model was used as a species tree prior. We ran PartitionFinder 2 (Lanfear, Frandsen, Wright, Senfeld, & Calcott, 2016) to identify the best nucleotide substitution model based on Bayesian information criterion (BIC): TRN+I+G for *COI* and *D-loop*, HKY+G for *Hodz3* and HAM-*ZBTB10-3*, HKY+G+I for *RTN4*, and K80+G for *RAG1.* For the six partitions, we assumed a lognormal relaxed clock (Drummond, Ho, Phillips, & Rambaut, 2006) using a lognormal distribution model with log mean of 0.01 and log standard deviation of 1. We ran five independent runs (15 ×10^7^ generations), sampling the parameters values and trees every 10,000 generations (Supporting file 1).

To time-calibrate the tree, we overcame the absence of *Hypostomus* fossils by applying an approach based on biogeographic calibrations. Ho and colleagues (Ho et al., 2015) listed two key assumptions for the application of such calibrations: i) a significant impact on population or species, and ii) available information on the age. We used four dated hydrogeological changes as calibration points that meet these assumptions: (1) Uplift of the Cordillera de Mérida (mean age of 8.0 Ma ± 0.08 at the crown age of *H. hondae + H. plecostomoides*); (2) Uplift of the Ecuadorian Andes (10 Ma ± 0.07 at the crown age of *Aphanotorulus* + *Isorineloricaria*); (3) Disconnection of the Tocantins-Araguaia River from the Amazon Basin (2.6 Ma ± 1 at the crown age of *Hypostomus* sp. ‘gr. *cochliodon*-Tar’ + *Hypostomus.* sp. ‘gr. *cochliodon*-Xin2’) (4) Boundary displacement between the Amazon and the Rio Paraguay systems (1 Ma ± 0.6 at the crown age of *Hypostomus* sp. ‘Rio Grande’ + *Hypostomus* sp. cf. *borelli,*). For more details regarding the calibration points, refer to the Supporting file 3. The inclusion of multiple calibrations may reduce the influence of eventual erroneous calibration and improve the robustness of the time tree (Duchêne, Lanfear, & Ho, 2014).

To evaluate the effect of the use of the aforementioned biogeographic events as calibration points in the *Hypostomus* time-tree, we performed three additional inferences (Supporting file 1). In each of these new reconstructions, one of the new calibration points was not used. We then assessed if important disparities in the time estimates among the distinct reconstruction are observed. Furthermore, assuming that the use of calibration points during phylogenetic reconstructions may improve accuracy of the phylogenetic reconstructions (Drummond et al., 2006), we also verified if major changes in the topology of the tree were observed. For these alternative time-tree reconstructions, we used the same parameters and priors as described before.

### Ancestral distribution range

To infer the biogeographic history of *Hypostomus*, we estimated the ancestral distribution range of all the nodes of the phylogenetic tree using the Dispersal–Extinction– Cladogenesis (DEC) model (Ree et al., 2005; Ree & Smith, 2008) implemented in RevBayes (Höhna et al., 2016) (Supporting file 2). The DEC model was originally described — and has been often used — as a maximum likelihood approach (Ree et al., 2005; Ree & Smith, 2008), but RevBayes offers a Bayesian implementation, It allows one to assess parameter uncertainty and to calculate Bayes Factor to compare competitive hypotheses (Landis, Freyman, & Baldwin, 2018).

Discrete biohydrogeographic regions (BHG) were defined based on the areas proposed by Abell et al. (2008), and on the areas of Neotropical freshwater fish endemism (Hubert & Renno, 2006; Montoya-Burgos, 2003; Vari, 1988). These areas were adapted to fit with the BHG regions that remained unfragmented during the historical changes in basin connectivity from the Miocene to the present. In this way, we defined 12 BHG regions (Figure 7A): (BHG region A) *Western Amazon* covers the upper portion of the Amazon Basin to the Middle Amazon River, including the sub-basins of Madeira and Negro rivers; (B) *Eastern Amazon* includes the lower portion of the Amazon River and tributaries draining in the Guiana and Brazilian shields, such as the Tapajós, Xingu and Trombetas rivers; (C) *Tocantins-Araguaia* River Basin; (D) *Orinoco* River Basin and the island of Trinidad; (E) *Guianas–Essequibo* includes the Guianese coastal rivers draining the Guyana Shield and flowing into the Atlantic Ocean; (F) *Paraguay* River System covers the Pilcomayo and Paraguay rivers and the lower section of the Paraná River; (G) *Upper Paraná* River covers the upper portion of the Paraná River; (H) *Uruguay* River Basin and *Lagoa dos Patos* System; (I) *Coastal Atlantic* drainages of the South- and Northeastern Brazil from the south of Santa Catarina state in Brazil to the, but not including, São Francisco River mouth; (J) *São Francisco* River Basin; (K) *Coastal Atlantic* drainages of Northern Brazil includes rivers flowing into the Atlantic Ocean between the Tocantins-Araguaia and São Francisco mouths; (L) trans-*Andean* drainages includes the Lake Maracaibo, Magdalena River Basin (which drain into the Caribbean Sea) and Pacific drainages of Ecuador and Colombia.

We defined two stationary biogeographic models, with constant constraints in dispersal possibilities for the complete time period of our study (25 Ma to present). The first model (M1) assumes full permeability between all pairs of BHG regions (connectivity value = 1), irrespective of whether they are adjacent to each other or not. The second model (M2) is semi-permeable, in which dispersion is possible but constrained between adjacent BHG regions (connectivity value = 0.1), and dispersion is not allowed between non-adjacent BHG regions (connectivity value = 0). One of the advantages of the DEC model is the possibility of combining *a priori* different temporal connectivity patterns among BHG regions by using a pairwise matrix of connectivity. Hence, we defined a third time-stratified model (M3) by taking into account historical changes in connectivity across BHG regions. For this aim, we used the following connectivity values: (i) a value of 1 was given to a pair of BHG regions for the period of time in which they displayed a full hydrological continuity (i.e. they were parts of a same basin); (ii) a value of 0.5 to a pair of BHG regions for the period of time in which they displayed partial hydrological continuity, for example through a permanent local connection such as the Casiquiare River currently connecting the Western Amazon with the Orinoco basins; (iii) a value of 0.1 to a pair of adjacent BHG regions showing no particular long standing water connection; and (iv) 0 to non adjacent BHG regions. We established these rules because fish, being limited to the aquatic environment, cannot disperse from non-adjacent regions in a single step. The M3 considered four time windows (Supporting file 3, Table S2), each one with a determined connectivity matrix according to the known paleo-landscape changes. For details, see Supporting file 3.

To run the time-stratified (M3) and stationary models (M1 and M2), we estimated a single average value of allopatry and subset sympatry rates. Moreover, the maximum ancestral range was set to three BHG regions, which corresponds to the maximum number of regions occupied by extant *Hypostomus* species included in our study. The original functions in RevBayes were originally designed to explore all the combinations of discrete BHG regions (Höhna et al., 2016). In our case, with a total of 12 BHG regions and setting the limit of ancestral areas to combinations of at most three BHG regions, we would have 298 possible combinations to compute. To optimize the analyses and reduce computation time, we modified the RevBayes script to disregard combinations of areas that have no biological meaning in terms of species distribution range, such as very distant areas with no present or past connectivity (*e.g. ABK*, *AGH* or *CDE*), reducing the number of possible ranges to 67. This new feature was key to this study since the biogeographic analyses defined using default RevBayes functionalities could not be successfully run for our dataset due to their excessive computational cost (i.e., requiring more than 32GB of RAM and several minutes per iteration). With our modification, the analyses did not suffer from memory limitations and ran faster (i.e., 17 seconds per iteration). A version of the helper script for biogeographic reconstructions in RevBayes can be found in the Supporting file 2 as well as in the GIT repository https://bitbucket.org/XavMeyer/biogeographyrevscript/src/master/, where improvements may be available. The results obtained with the different biogeographic models were compared and ranked by calculating the Bayes Factor, which is a typical model selection approach in the Bayesian framework (Baele, Lemey, & Vansteelandt, 2013). To do so, we ran 40’000 iterations sampling every 10 iterations. Convergence was assessed by plotting log parameters in Tracer.

### Habitat preference evolution

In order to investigate how ancestral ecology played a role on the biogeographic history of *Hypostomus*, we evaluated the pattern of evolution of the habitat preference across the phylogenetic tree of *Hypostomus*. We reconstructed the ancestral habitat preference by defining the nine categories of habitats that were modified from those according to (2011), as listed below (Figure 6):

1. *Slow flowing small streams* (SFSS): streams with maximum 50 m width, such as headwater streams or small tributaries, being localized either in uplands or in lowlands, generally muddy, with substrate comprising mostly of clay and sand.
2. *Medium to fast flowing small streams* (FFSS): as the previous category, but substrate comprised mainly of rocks or small stones.
3. *Slow flowing medium-sized rivers* (SFMR): rivers with *c.* 50–1,000 m width, either placed in uplands or lowlands, with muddy and/or sandy substrate.
4. *Medium to fast flowing medium-sized rivers* (FFMR): as in SFMR, but substrate mainly composed of gravel, stones and rocks. Medium-sized rapids and waterfalls are included in this category.
5. *Slow flowing large rivers* (SFLR): large rivers with more than 1 km width, which include all the rivers or portions of them with very slow flowing waters, with substrate composed mainly of clay and sand.
6. *Medium to fast flowing large rivers* (FFLR): as in SFLR, but with gravel, rocky and stony substrates, including large rapids and waterfalls.
7. *Floodplains and ponds* (FLPO): areas that are seasonally inundated by the main rivers, including *várzea* and *igapó* lakes, ponds, flooded forests, flooded savannahs and flooded grasslands.
8. *Freshwater estuarine systems* (FRES): areas of estuary of very large rivers, where the salinity is still very low. In this category we included only the Río de La Plata, comprising specifically the mouth of Uruguay and Paraguay rivers.
9. *Brackish water* (BRWA): estuary and coastal regions characterized by relatively high salinity. A single *Hypostomus* species lives in this very atypical habitat in the coast of the Guyanas.

We used BEAST 1.8.1 (Drummond, Suchard, Xie, & Rambaut, 2012b) to reconstruct the ancestral habitat state (Supporting file 1). We employed the aforementioned time-calibrated tree, by fixing topology and branch lengths. We assumed a symmetric model for discrete state reconstructions. A strict clock was used, thereby enforcing a homogeneous rate of trait evolution across all branches, and the prior on mean rate of habitat evolution was set as a gamma distribution. Three independent runs of 5 × 10^7^ generations, sampled every 5,000^th^ generation, were ran. Convergence of the runs and ESS >200 were confirmed with Tracer 1.6. The post-burn-in samples from the three runs were combined using LogCombiner 2.4.8 to produce a consensus tree in TreeAnnotator 2.4.8.

## Results

### A robust calibrated Phylogeny of Hypostomus

We inferred a time-calibrated phylogeny of *Hypostomus* using four calibration points (calibration scenario 1) indicating that the *Hypostomus* lineage trace back to 14.7 Ma (HPD 95% = 17.8–11.4 Ma), which corresponds to the split between *Hypostomus* and its closest relatives *Hemiancistrus fuliginosus + Hemiancistrus punctulatus* (Figure 1). The estimated age of the Most Recent Common Ancestor (MRCA) of *Hypostomus* is 12.1 Ma (14.3–10.1 Ma) while the ages of divergence between the main *Hypostomus* lineages are presented in our calibrated phylogeny (Figure 1).

**Figure 1.**
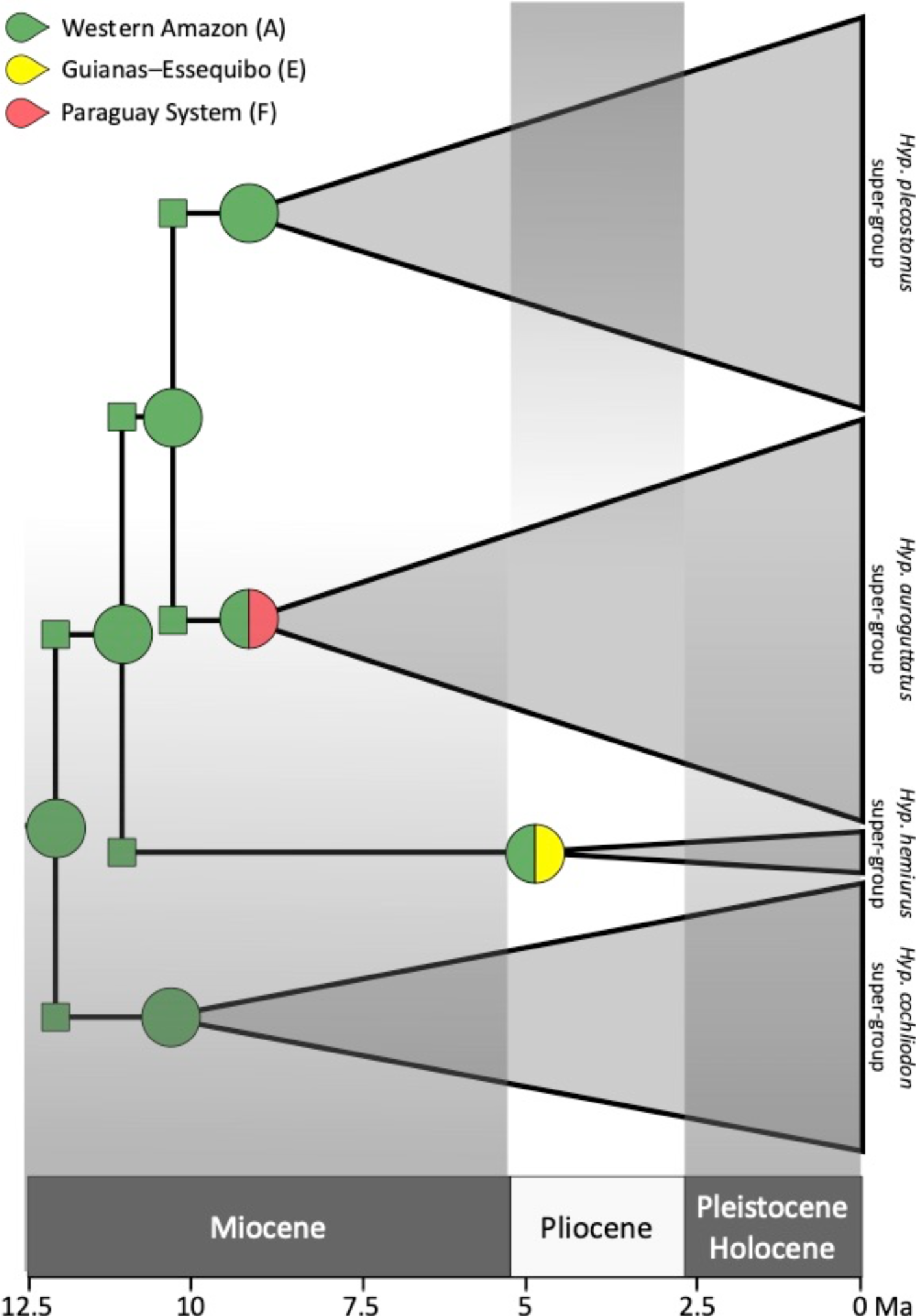
Ancestral range reconstruction of the first ancestral *Hypostomus*and of the ancestors of the four main internal lineages. For detailed results, see Figs. 2–5. The posterior probabilities of the ancestral range reconstructions shown here are ≥ 0.55.

To evaluate the robustness of our calibrated phylogeny, we assessed the compatibility of each calibration point with regards to the three others by running three new calibrated phylogenetic inferences with the exclusion of one of the four calibrations at a time (calibration scenarios 2 to 4 in Table 1). The resulting topologies were very similar among each other and also with the results obtained with the full set of calibration points (Supporting file 1). The super-groups of *Hypostomus*, as previously defined in the literature (Jardim de Queiroz et al., 2020), where reconstructed with strong posterior probability (PP ≥ 0.97) in all calibration scenarios (Supporting file 1). The only noticeable difference was the age retrieved for the calibration point 2 consisting in the split between *Isorineloricaria* and *Aphanotorulus* (IA split) attributed to the uplift of the Ecuadorian Andes. When this calibration point was not used as a prior (calibration scenario 4), the age of this split was estimated to be 19.6 Ma (27.3–13 Ma), that is, older than the uplift of the Ecuadorian Andes, the hypothesized vicariant event causing the split. We also noticed that the splitting ages at the deepest nodes were in general more ancient when the IA split was not considered as a calibration point (calibration scenario 4). However, in all these cases, there is an important overlap of the HPD 95% across the different calibration scenarios (Table 1). Therefore, we used the scenario with four calibration points (calibration scenario 1) in the downstream analyses.

**Table 1.**
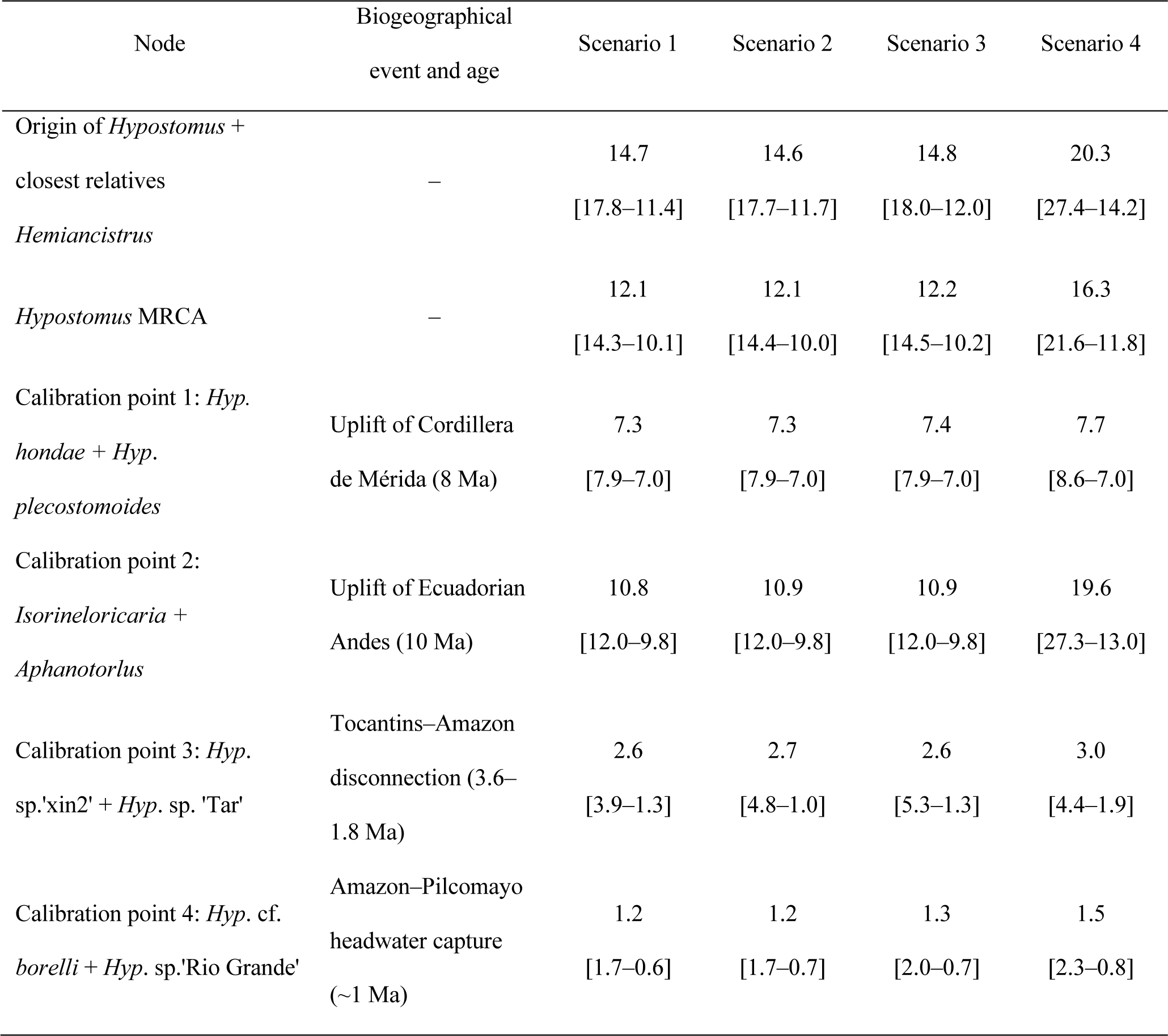
Retrieved ages and Highest Posterior Density (HPD) 95% (between brackets) of six representative nodes of the *Hypostomus* phylogenetic tree. Four calibration analyses were performed (Supporting file 1). In *scenario 1*, from which the results of this work are based on, all four-calibration points were used as priors; in *scenario 2*, the calibration point 3 (Tocantins–Amazon disconnection) was not included; in *scenario 3*, the calibration point 4 (Amazon–Pilcomayo headwater capture) was not included; and in *scenario 4*, calibration point 2 (Uplift of Ecuadorian Andes) was not included. MRCA: most recent common ancestor.

### Western Amazon was a centre of origin and dispersal

We reconstructed the biogeographic history of the genus *Hypostomus* using three biogeographic models in RevBayes. The first two were stationary models in which the connectivity between biohydrogeographic (BHG) regions remained unchained along time: the first with full permeability between all pairs of BHG regions (Model 1, M1), and the second combining (i) limited permeability between adjacent BHG regions and (ii) no permeability between non-adjacent ones (Model 2, M2). The third model (M3) assumed a time-stratified scenario with connectivity patterns adapted to four time windows, which were defined based on the most important hydrological changes documented for South America. The time-stratified model M3 showed the highest score relative to the two stationary models (Bayes Factor difference > 19; Supporting file 3, Table S1), providing the best approximation of ancestral ranges throughout the radiation of *Hypostomus,* which we used in our further analyses. Accordingly, the most recent common ancestor (MRCA) of the lineage comprising *Hypostomus* and its sister *Hemiancistrus* group lived in the Western Amazon + Paraguay (start state = region AF; posterior probability = 0.44; Supporting file 1a). This ancestor then experienced a vicariant event (AF → A|F), by which the ancestral population isolated in the Western Amazon (A) gave origin to the first ancestral *Hypostomus*. While living in Western Amazon, this ancestral *Hypostomus* split into consecutive lineages being at the origin of the four recognized super-groups of *Hypostomus* (*Hyp. cochliodon* super-group, *Hyp. hemiurus* super-group, *Hyp. auroguttatus* super-group and *Hyp. plecostomus* super-group). The divergences among these super-groups followed a pattern where one of the descendants remained in Western Amazon, while its sister descendent colonized a new BHG region (Figure 1). In this way, a population from Western Amazon colonized the Guianas–Essequibo region (A→AE) giving rise to the ancestor of the *Hyp. hemiurus* super-group (HHsg). Subsequently, stemming from the ancestral species that remained in Western Amazon, a population entered into the Paraguay System (A→AF), initiating the *Hyp*. *nematopterus* + *Hyp. auroguttatus* super-group (HAsg). Finally, the ancestral species still residing in Western Amazon gave rise to the *Hyp*. *plecostomus* super-group (HPsg).

Similarly, other Neotropical main basins where gradually colonized by *Hypostomus* representatives stemming from Western Amazon. The first colonization of the Orinoco Basin (BHG region D) took place around 10.5 Ma with the arrival from Western Amazon (A→D) of the ancestor of *Hyp*. *sculpodon* (a member of the HCsg lineage; Figure 2). The Lower Amazon Basin (Eastern Amazon, BHG region B) was also colonized first by emigrants stemming from Western Amazon at 9.9 Ma (A→BC; Figure 2). The *trans*-Andean area (BHG region L) was the only region adjacent to the Western Amazon that was not initially colonized by Western Amazon ancestors, but by *Hypostomus* stemming from the Orinoco Basin (D → DL; Figure 2).

**Figure 2.**
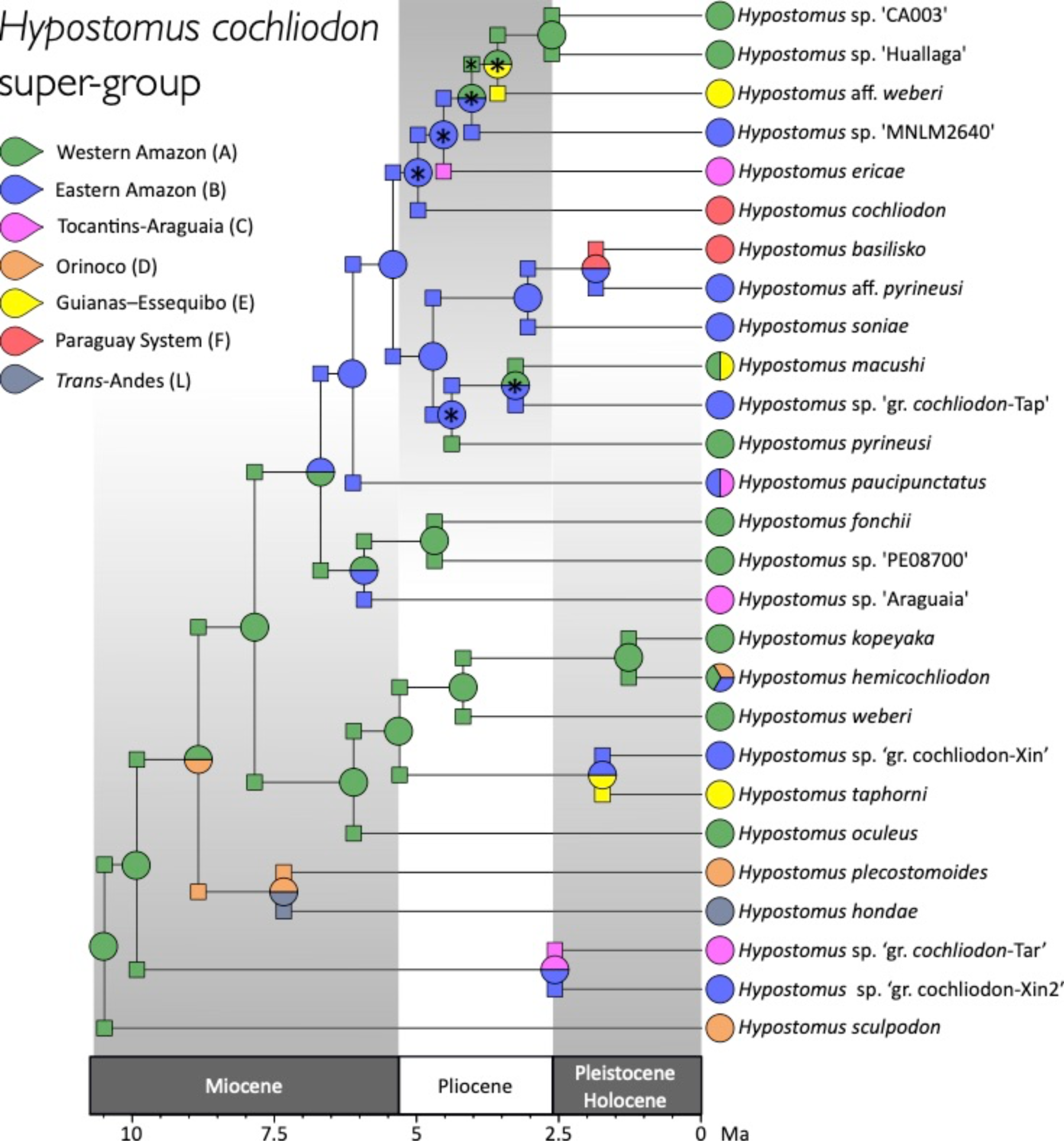
Ancestral range reconstruction of the *Hypostomus cochliodon* super-group. (* posterior probability of ancestral range reconstruction < 0.5).

According to our biogeographic reconstructions, the Lower Amazon Basin (BHG region B), the Orinoco Basin (BHG region D), and the Guianas-Essequibo System (BHG region E) were repeatedly colonized by *Hypostomus* species coming from Western Amazon. For example, the Orinoco Basin was independently colonized by Western Amazon ancestors at least in three instances: once in the lineage leading to *Hyp*. *hemicochliodon* (A→ABD; 1.3 Ma, Figure 2), once in the lineage leading to *Hyp*. *hondae* + *Hyp*. *plecostomoides* (A→AD; 8.8 Ma, Figure 2), and once at the root of the *Hyp. robinii* group (A→AD; 7.1 Ma, Figure 5). The Guianas-Essequibo System, besides the early colonization by the ancestor of the HHsg, was independently colonized two additional times by Western Amazon ancestors, the first leading to the monotypic lineage of *Hyp*. *nematopterus* (A→E; 9.2 Ma, Figure 4) and the second giving rise to the *Hyp. watwata* group (A→AE; 7.3 Ma, Figure 5). The re-colonization of the Lower Amazon by Western Amazon emigrants can be exemplified by the arrival of the ancestor of today’s *Hyp.* sp. ‘Paru’ (A→AB; 5 Ma, Figure 3).

**Figure 3.**
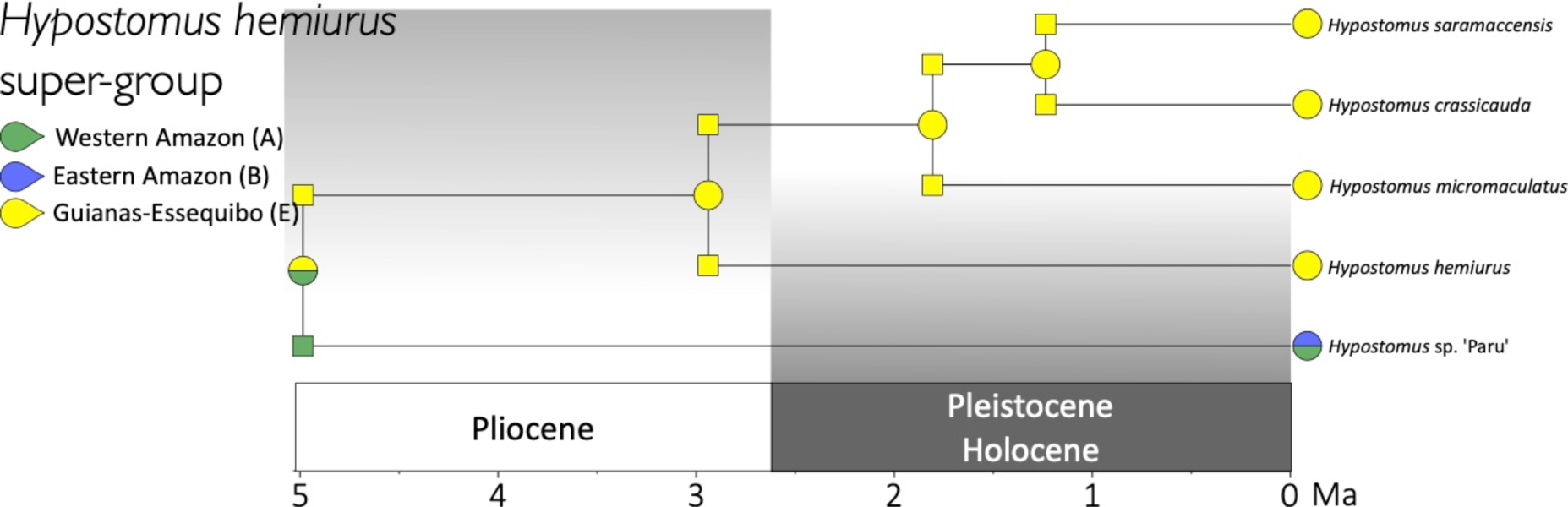
Ancestral range reconstruction of the *Hypostomus hemiurus* super-group. The posterior probabilities of ancestral range reconstructions shown here are ≥ 0.54).

**Figure 4.**
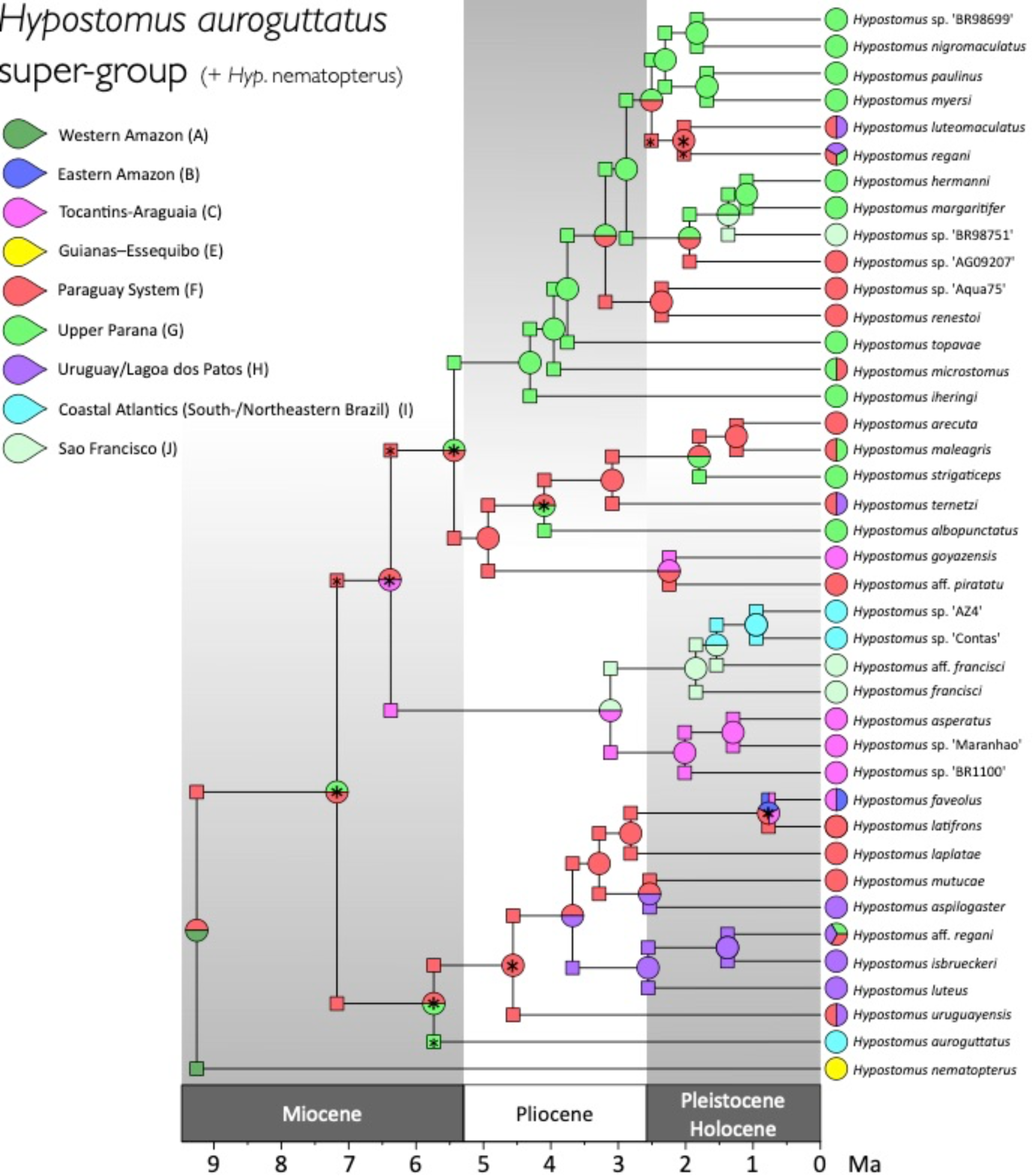
Ancestral range reconstruction of the clade that contains the *Hypostomus auroguttatus* super-group and *Hypostomus nematopterus*. (* posterior probability of ancestral range reconstruction < 0.5).

**Figure 5.**
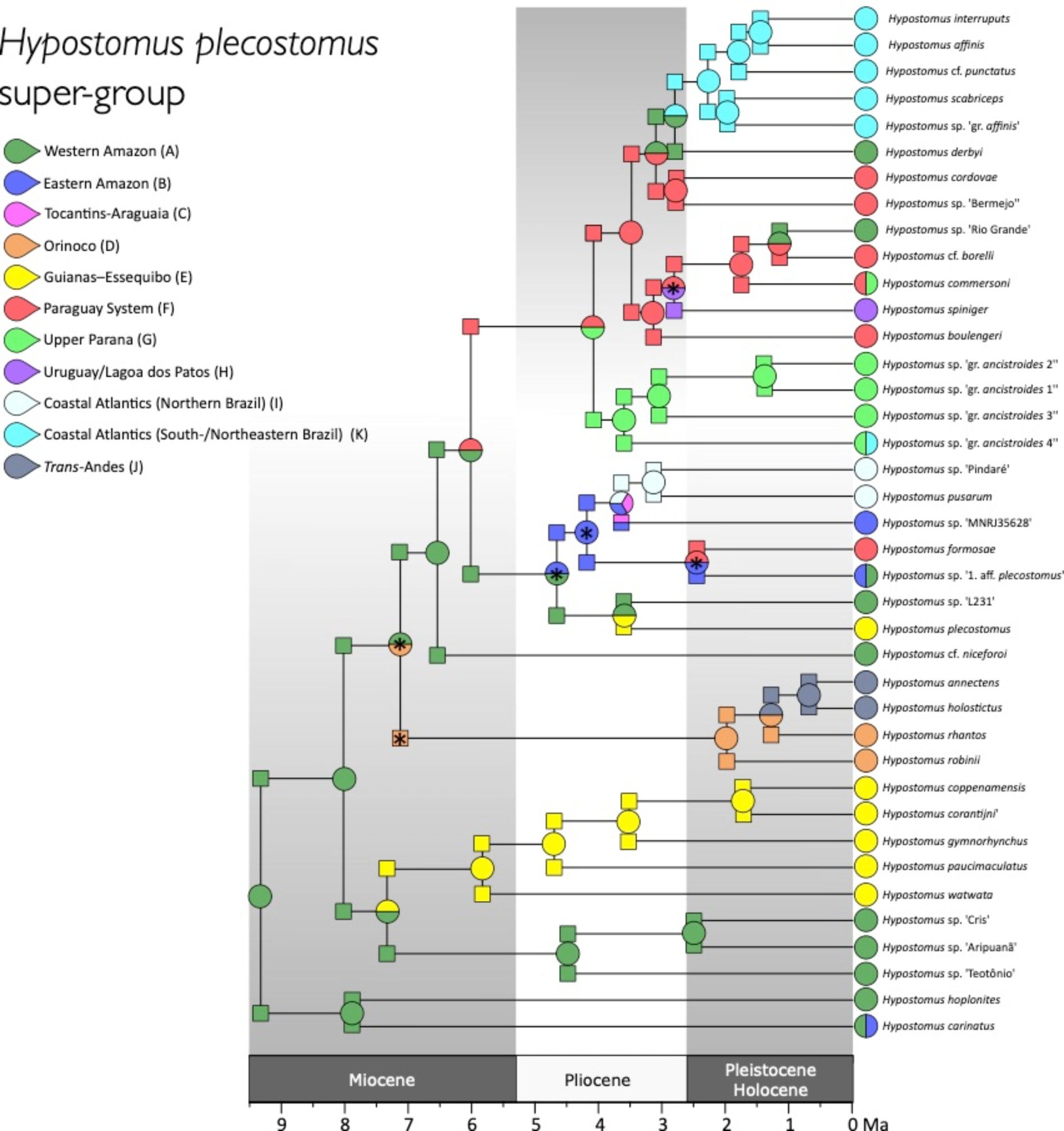
Ancestral range reconstruction of the *Hypostomus plecostomus* super-group. (* posterior probability of ancestral range reconstruction < 0.5).

**Figure 6.**
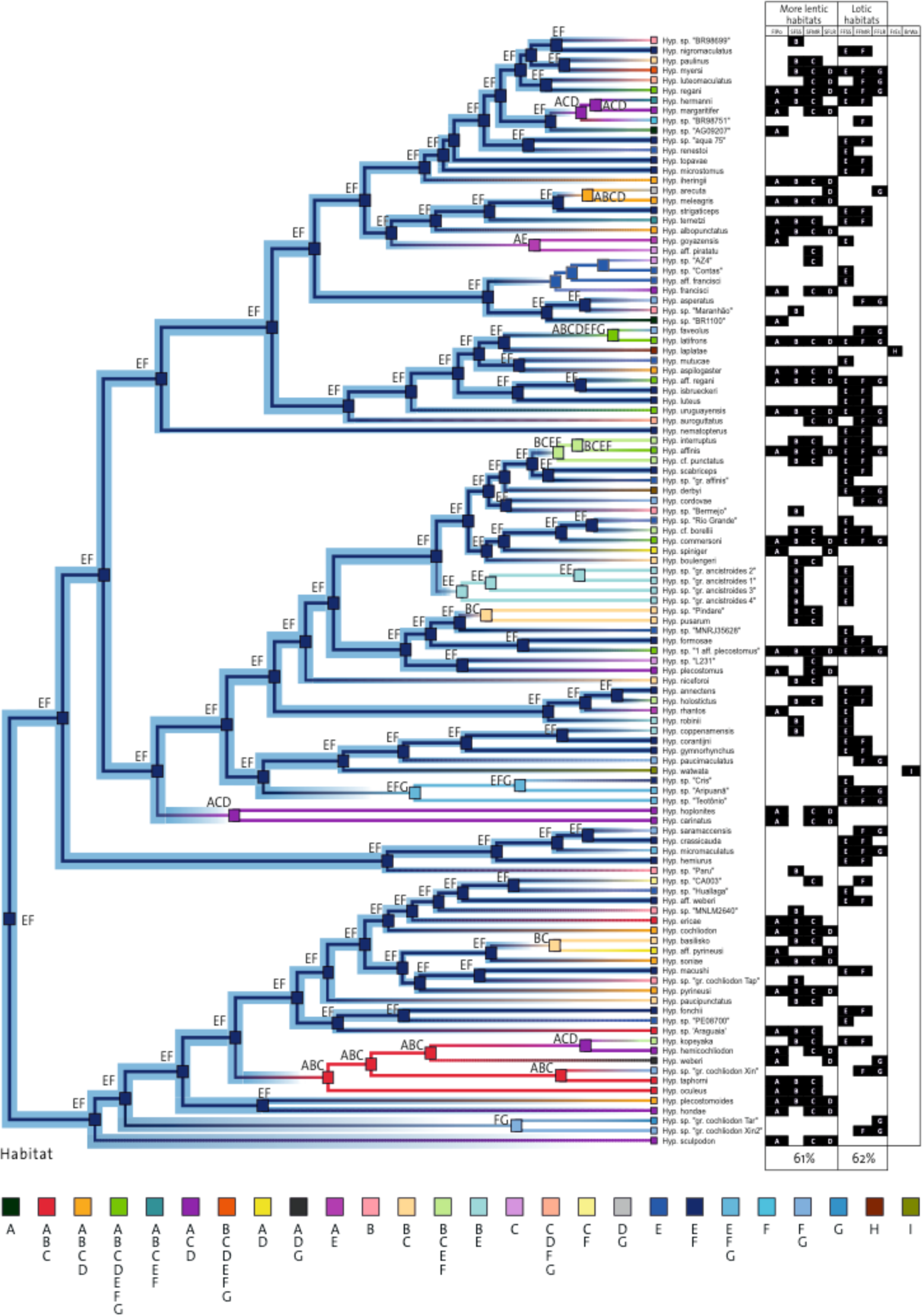
Ancestral habitat reconstruction of *Hypostomus*. A = floodplains and ponds (FlPo); B = slow flowing small streams (SFSS); C = slow flowing medium rivers (SFMR); D = Slow flowing large rivers (SFLR); E = medium to fast flowing small stream (FFSS); F = medium to fast flowing medium rivers (FFMR); G = medium to fast flowing large rivers (FFLR); H = freshwater estuarine systems (FrEs); I = brackish water (BrWa). More information, including habitat reconstruction of outgroups and posterior probabilities can be found in Supporting file 1.

**Figure 7.**
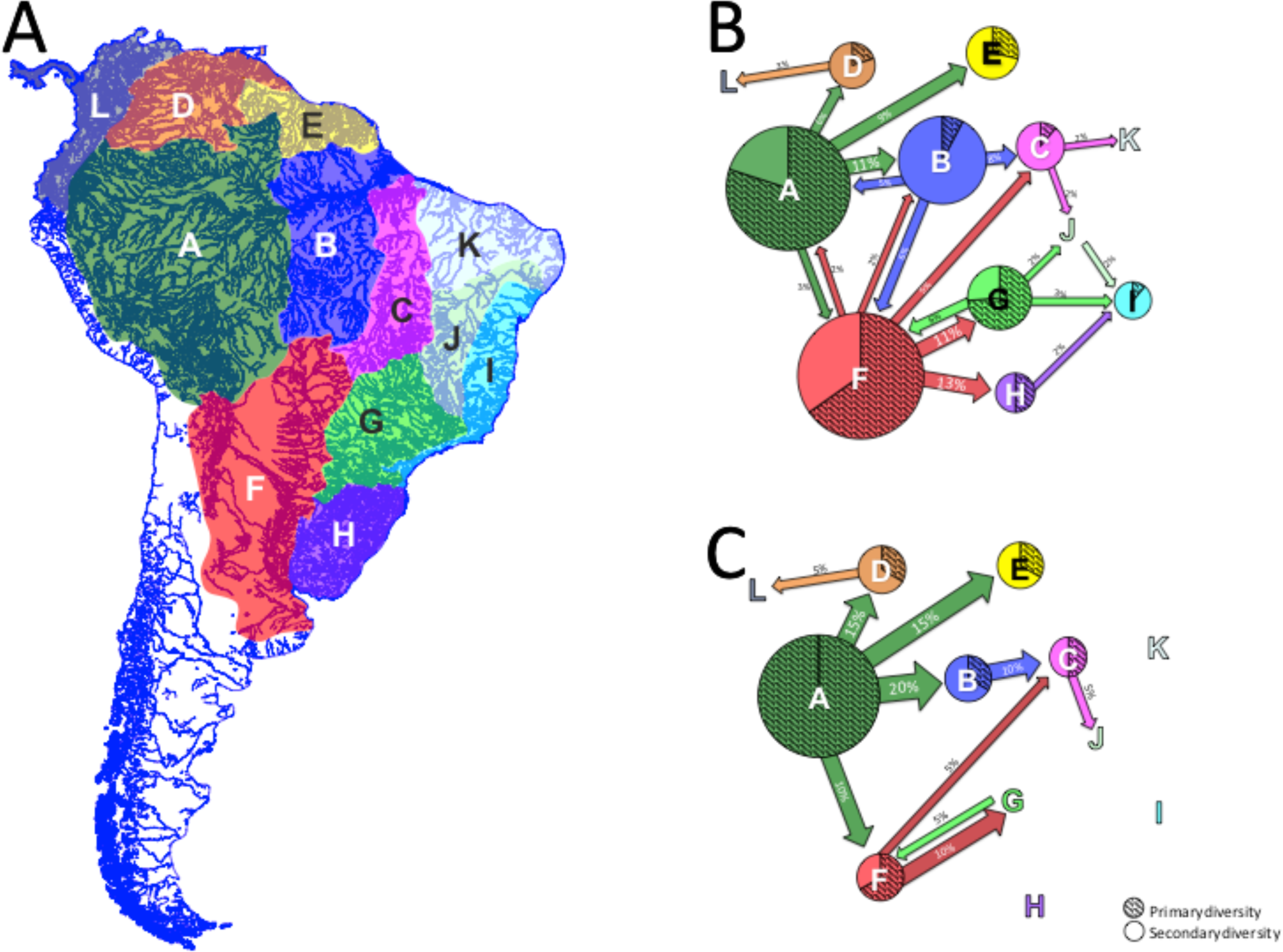
Dispersal events between biohydrogeographic regions (BHG). A) Map showing the division of South America into BHG regions. B) Dispersal events inferred from 12.1 Ma to the present. C) Dispersal events inferred from 12.1 to 6 Ma. Circle sizes are proportional to the number of extant *Hypostomus* species. Dashed areas of the circles represent “primary diversity”, i.e. the proportion of *Hypostomus* species that originated from the first colonization of the region. Species that arose from secondary colonization events were considered as “secondary diversity”. As the Western Amazon (region A) was the place where *Hypostomus* originated, we considered as primary diversity only the species that showed no ancestor distributed in another BHG region.

### Hypostomus radiation into Southern and Eastern South American basins

The colonization of the Upper Paraná (region G) started with the ancestor of the HAsg at 9.2 Ma (Figure 4), stemming from the pre-existing Paraguayan lineage (F→FG). Subsequently, also within the HAsg, the first colonization of the Uruguay System/Lagoa dos Patos (region H) took place at about 4.6 Ma, with colonists emigrating from the Paraguay System (F→FH, Figure 4). Moreover, we recovered two independent colonization of the São Francisco River (region J), both within the *Hyp. auroguttatus* super-group (HAsg). The first gave origin to the *Hyp. asperatus* group (C→CJ; 6.4 Ma,), while the second was the ancestor of todays *Hyp.* sp. ‘BR98751’ (G→GJ; 1.9–1.4 Ma), a member of the *Hyp. regani* group (Figure 4).

A single colonization of the coastal Atlantic drainages of Northern Brazil (region K) was inferred and involved the ancestor of *Hyp*. *pusarum* + *Hyp*. sp. “Pindaré” (B→BCK; 4.2 Ma, Figure 5). On the other hand, the coastal Atlantic rivers (region I) were colonized four independent times. In two cases the ancestral colonists stemmed from the Upper Paraná: (i) the ancestor of *Hyp. auroguttatus* (belonging to the HAsg) at 5.7 Ma (G→I, Figure 4) and (ii) the ancestor of *Hyp*. sp. “gr. *ancistroides* 4” (HPsg) (G→GI) at 3.6 Ma (Figure 5); while in the third case (iii) the colonists came from a region composed of the Paraguay plus the Upper Paraná, at about 3.2 Ma, which led to a small monophyletic lineage comprising five species of our dataset, including *Hyp. affinis* and relatives (FG → GI, Figure 5); and in the fourth case (iv) the ancestor stemmed from the São Francisco Basin at around 1.8 Ma and led to the group of species *Hyp*. sp. “AZ4” + *Hyp*. sp. “Contas” + *Hyp*. aff. *francisci* (J→JI, Figure 4).

### Habitat preference conservatism with recent shifts

The ancestral habitat preference reconstructed with BEAST revealed that the MRCA of *Hypostomus* was living in small to medium size streams and rivers with medium–fast flowing waters (EF, Figure 6). This type of habitat remained the preferred one along most of the evolutionary history of the genus and still persists in many representatives of all main lineages. This result is indicative of marked niche conservatism in this genus. Among the changes in habitat preference, the most ancient shift is observed in the stem branch of the lineage including *Hyp. hoplonites* + *Hyp. carinatus* about 7.9 Ma, as this lineage adapted to more lentic environments (floodplains and slow flow medium-large rivers, ACD). The next shift in habitat preference occurred at about 6.1 Ma within the HCsg lineage, with a change towards more lentic environments (floodplains + slow flow medium–large rivers, ACD; and floodplains + slow flow small–medium rivers, ABC). All the remaining shifts in habitat preference occurred quite recently, starting at about 4.5 Ma. Today, the proportion of species inhabiting lentic habitats is approximately equal to the proportion living in lotic habitats (Figure 6).

## Discussion

Although the Amazon region has been recognized as the major source of diversity of terrestrial organisms in the Neotropical realm [e.g. 15, 16], an equivalent role for freshwater lineages has not been thoroughly investigated. In the present study, we have addressed this issue by analyzing *Hypostomus* as a model group due to its remarkable species-richness, its continental-wide distribution, and because its species have colonized all Neotropical freshwater habitats. We assembled information about the distribution, ecology and habitat preference of 111 *Hypostomus* species from the main basins of South America and we calibrated a large phylogenetic tree. We also took advantage of the growing knowledge related to the history of river basins and landscape changes in Tropical South America (J. S. Albert & Carvalho, 2011; J. S. Albert & Reis, 2011; Carvalho & Albert, 2011; e.g. C. Hoorn et al., 2010; John G. Lundberg, Pérez, Dahdul, & Aguilera, 2011; Tagliacollo, Roxo, Duke-Sylvester, Oliveira, & Albert, 2015) to define time windows for each configuration of river network and connectivity, leading to more realistic biogeographic reconstructions. To improve our biogeographic reconstructions, we developed a new tool for RevBayes allowing for computationally tractable analyses with more precise geographic divisions by exploiting *a priori* biologically meaningful distribution ranges.

### Western Amazon was the centre of origin of Hypostomus

Deciphering the precise distribution range of a distant ancestor based on its current day descendants depends largely on the completeness of the species sampling, the knowledge of the species distribution range and the size and number of the geographic areas considered in the analysis. Previous attempts to infer the ancestral range of *Hypostomus* used restricted taxonomic sampling, and resulted in excessively large ancestral distribution range for the first ancestral *Hypostomus*, encompassing the entire proto Amazon-Orinoco System (Cardoso et al., 2021) or even the proto-Amazon-Orinoco plus Upper Paraná (Silva et al., 2016). With our comprehensive taxonomic sampling of *Hypostomus* and close relatives, we found that the MRCA of *Hypostomus* inhabited the Western Amazon, an area that nowadays corresponds to the Upper Amazon Basin including some important tributaries, such as the Madeira, Negro, Purus and Japurá rivers. Our calibrated phylogeny based on four cross-tested calibration points (Figs. 1–5; Supporting file 1: Time tree 1) indicates that the first ancestral *Hypostomus* emerged in the Middle-Miocene, approximately from 14.7 Ma, and started to diversify at approximately 12.1 Ma. These temporal results are in agreement with previous hypotheses (J. I. Montoya-Burgos, 2003; Silva et al., 2016), but contrast with the Oligo-Miocene origin (∼25 Ma) proposed recently (Cardoso et al., 2021).

Our findings indicate that the first ancestral *Hypostomus* originated and lived in the Western Amazon Region, at a time where this region was mostly occupied by the Lake Pebas. This mega basin was composed of a mosaic of lentic water bodies and wetlands that dominated the lowlands of western Amazonia, bordered by the slopes of the Andes on the western side and of the Brazilian and Guyana shields on the eastern side (C. Hoorn et al., 2010; Lovejoy, Bermingham, & Martin, 1998; Wesselingh et al., 2006).

Due to marine transgressions, the lentic waters of the Lake Pebas basin are suggested to have been influenced by the sea, increasing its salinity (Lovejoy et al., 1998; Wesselingh et al., 2006). It is unlikely that the ancestral *Hypostomus* lived in this brackish-like lentic habitat, as we found that adaptation to brackish waters occurred only once and recently in the evolutionary history of *Hypostomus*, with *Hypostomus watwata* inhabiting the estuaries of Guianese rivers. Moreover, our habitat preference reconstructions indicate that the ancestral *Hypostomus* was adapted to small rivers with medium-to-fast flowing waters. It is therefore most likely that the first *Hypostomus* was distributed in small rivers draining the slopes of the mountains surrounding Lake Pebas, either on the Andean side or on the slopes of the Purus Arch.

According to our biogeographic analysis, the Western Amazon is the main biohydrogeographic region from which most of the dispersal events originated during the early radiation of *Hypostomus* (Figure 7). In total, the Western Amazon accounted for 30% of all the reconstructed dispersal events along the evolutionary history of *Hypostomus*, a proportion only reached by the Paraguay System (∼31%). However, when we consider only the first half of the *Hypostomus* radiation period (12.1–6 Ma), the Western Amazon was the dominant center of dispersal, accounting for 60% of these events.

We uncovered that Western Amazon was the center of origin of *Hypostomus*, a result that is in agreement with the few available findings on Neotropical fish. In a recent study of Amazonian fish biodiversity, the Western Amazon area has been suggested to be the main geographic region of origination of the Amazonian fish fauna, with a downstream colonization of the lower Amazon basin (Oberdorff et al., 2019). However, this study was geographically restricted to the Amazon basin only. At a wider spatial scale, detailed fish biogeographic analyses revealed that the entire Amazon basin, taken as a single BHG region, was the center of origin and dispersal into other Neotropical basins for the catfish subfamily Hypoptopomatinae (Chiachio, Oliveira, & Montoya-Burgos, 2008) and for the characiform genus *Triportheus* (Mariguela et al., 2016). Recently, a more refined biogeographic reconstruction also revealed the Western Amazon as the origin of marine-derived Neotropical freshwater stingrays (Fontenelle et al., 2021), yet the marine origin of this group may not be representative of other primary freshwater lineages. Therefore, our results bring new evidence in support of the hypothesis that Western Amazon was a primary center of freshwater fish origination.

### The role of ecology in Hypostomus dispersal

The preferred ancestral habitat of *Hypostomus* consisted in small-to-medium streams with medium-to-fast flowing waters (Figure 6). This habitat preference predominated in the early evolution of *Hypostomus*, while the colonization of new habitats, such as lakes, floodplains, large rivers and estuaries occurred more recently. According to this eco-evolutionary hypothesis, the first half of the *Hypostomus* radiation was characterized by a strong niche conservatism, which may have played a role on the dispersal and speciation modes of this genus. The preferred ancestral habitat of *Hypostomus* is very common in river headwaters, which are portions of drainages prone to be captured by adjacent basins through evolutionary time. Indeed, the geomorphological history of South America has been marked by many events of headwater captures (James S. Albert et al., 2018; Lavarini, Magalhães Júnior, Oliveira, & Carvalho, 2016; Stokes, Goldberg, & Perron, 2018), their incidence being increased by the active tectonics of this continent and by river meandering and erosion in relatively flat relief in several watershed divides.

Relying on our findings of *Hypostomus* ancestral habitat preference and strong niche conservatism along most of their early radiation, we propose that much of the dispersal events into new drainages occurred through the process of headwater captures, promoting range expansion and subsequent allopatric speciation due to geographic isolation. In support of this hypothesis, the results of the ancestral habitat reconstructions indicate that ∼61% of dispersal events along the evolution of *Hypostomus* took place in species preferring small or medium rivers with fast-flowing waters (lotic habitats). This is an expected outcome of dispersal by headwater captures, and could hardly be explained by dispersal through alternative environments, such as river delta connections during low sea-level periods (Cardoso & Montoya-Burgos, 2009; A. T. Thomaz, Malabarba, & Knowles, 2017), as the habitat preference of the dispersing species does not match the size of the water body or the stillness of the water found in such environments. The role of headwater captures in dispersal and speciation in South American freshwater fishes has been often suggested, and the hydrogeological hypothesis of diversification (J. I. Montoya-Burgos, 2003) was coined to incorporate this phenomenon. Evidence of headwater captures have been found in drainage boundaries all over Tropical South America, such as among Guianese rivers (e.g. Cardoso & Montoya-Burgos, 2009), between the Amazon Basin and Guianese rivers (Hubert & Renno, 2006), between the Amazon and the Orinoco basins (J. G. Lundberg et al., 1998), between the Amazon Basin and the Paraguay System (Hubert & Renno, 2006; J. G. Lundberg et al., 1998; J. I. Montoya-Burgos, 2003; Tagliacollo et al., 2015), between the Upper Paraná and the São Francisco river (Machado, Galetti, & Carnaval, 2018), among Atlantic coastal rivers (M. C. S. L. Malabarba, 1998; Ribeiro, 2006; Roxo et al., 2012), and among many others.

However, river headwaters are not always located in mountainous landscapes and temporary connections in basin divides located in flatlands might have also been dispersal routes. For instance, the headwaters of the Guaporé River (Amazon Basin) and Paraguay are located in a flat landscape (around 500 m in altitude) and are regularly interconnected during rainy seasons (Carvalho & Albert, 2011). However, this hypothesis fits better with dispersals later in the radiation of *Hypostomus*, when species shifted their habitat preference for more lentic environments.

Every dispersal event into a new basin might have been an ecological opportunity for *Hypostomus* to occupy its preferred habitat, with no congeneric competitors. Although more distant relatives were likely present in the newly colonized basins, we can speculate that a perfect niche overlap between the residing distant relatives and the new *Hypostomus* colonists is unlikely, although our data did not allow us to test this hypothesis. The hypothetical absence of strong competitors in basins colonized for the first time by *Hypostomus* would explain the marked niche conservatism characterizing the early evolutionary history of this genus, a hypothesis that can be tested in future studies.

We hypothesize that when basins were colonized for the second time, *Hypostomus* started to shift habitat preference, possibly due to competition between resident and colonist congeneric species. This hypothesis is supported by the finding that *Hypostomus* species started to diversify their habitat preference relatively recently along their evolutionary history, when the new dispersers where invading basins already occupied by congenerics. We suggest that the diversity in habitat preference among extant *Hypostomus* species (Figure 6) is most likely the outcome of growing intra-generic competition triggered by the repeated colonization of occupied rivers. According to radiation theory, niche shift is a response to inter-specific competition that allows the co-existence of closely related species, increasing local species-richness (Brew, 1982; Schluter, 2000). Consequently, our findings provide instrumental evidence for explaining the remarkable species richness, species coexistence and the wide distribution range of *Hypostomus* in the Neotropics. The eco-evolution of *Hypostomus* diversity, ecology and distribution may serve as evolutionary model for the entire fish community to which *Hypostomus* belongs, and our results may be informative for understanding the evolutionary history of fish taxa with comparable biological and ecological traits.

### Western Amazon as a fish center of dispersal

Elaborating on our findings, we propose that Western Amazon is a center of fish dispersal for five main reasons. (I) It is the center of origin of *Hypostomus*, and this is probably the case for many other species-rich Neotropical fish lineages. (II) Western Amazon is geographically located at the heart of Tropical South America, sharing drainage divides with almost all other main river basins with numerous episodic ichthyologic interchanges through headwater captures. (III) During the Miocene to the present, Western Amazon was the BHG region that had the largest amount of long lasting hydrological interchanged with neighboring basins (Supporting file 3, Table S2). (IV) Western Amazon encompasses the largest Neotropical freshwater system, and is the most fish species-rich BHG region nowadays and was already remarkably rich during the Neogene (John G. Lundberg et al., 2011). Together, these characteristics help to explain why Western Amazon exported repeatedly colonists into its neighboring biogeographic areas all along the evolutionary history of *Hypostomus*. We argue that these characteristics may also be valid for many other Tropical South American fish lineages that had Miocene representative in Western Amazon, as this region acted as a distribution platform, boosting their dispersion throughout Neotropical freshwaters.

### Conclusions

In the present work, we assessed the assumption that the Amazon basin was a major center of fish dispersal, spreading new species into neighboring Neotropical river basins from the Miocene to the present. Our detailed time-stratified biogeographic reconstruction of *Hypostomus*, one of the most species-rich genera in the Neotropical Region, indicated that this group emerged in the Western Amazon in the Middle Miocene, when the palaeolandscape was completely different from what we observe nowadays. While new *Hypostomus* species were gradually accumulating in Western Amazon, their ancestral habitat preference enabled them to colonize niches devoid of closely related competitors, and through headwater captures, multiplied the ecological opportunities to spread and diversify into newly colonized basins. Dispersal out of Western Amazon was also boosted by the reconfiguration of the paleo-Amazon-Orinoco watershed, with the disconnection of the Orinoco basin and the junction with Eastern Amazon. The central geographic location of Western Amazon, its longstanding connectivity with other basins, its large extension composed of many sub-basins hosting growing number of species, and the headwater captures spreading *Hypostomus* species in adjacent basins, altogether designate Western Amazon as a center of origin and dissemination for *Hypostomus*. As these characteristics might hold true for many fish lineages with Miocene representatives in Western Amazon, the pattern we observed here for *Hypostomus* may also be valid for a significant fraction of the Neotropical fish diversity. This scenario is supported by the fact that the Western Amazon is an extensive area located at the heart of Tropical South America, with long lasting hydrological connections with neighboring basins, as well as multiple episodic fish exchanges via headwater captures.

## Acknowledgments

We thank Margaret Zur and Mary Burridge from the Royal Ontario Museum (Canada) for sample donating (*Hypostomus annectens)*, Jaime Sarmiento and Soraya Barrera from the Museo Nacional de Historia Natural, La Paz, for sample sharing, and Mike R. May for RevBayes-related advises. Part of the analyses was performed on the Baobab cluster, a high performance-computing cluster of the University of Geneva. Funding were provided by *Conselho Nacional de Desenvolvimento Científico e Tecnológico*, Brazil (Program Sciences without Borders, CNPq 229237/2013-4, granted to LJQ; CNPq 423526/2018-9 and 312801/2017-3, granted to PAB; CNPQ 307775/2018-6 and 424668/2018-1, granted to TEP); Brazilian–Swiss Joint Research Programme, Switzerland (BSJRP S18794, granted to JIMB, GTV and LJQ); *Comissão de Aperfeiçoamento do Pessoal de Nível Superior,* Brazil (CNPq/CAPES 88882.156885/2016-01 granted to PAB); *Donation Claraz*, Switzerland (granted to JIMB); *Fundação Carlos Chagas Filho de Apoio à Pesquisa do Estado do Rio de Janeiro* (FAPERJ 200.063/2019 granted to PAB); Swiss Seed Money Grants Latin America 2015, Switzerland (granted to JIMB and YPC); Swiss National Science Foundation (SNSF 3100A0-104005, granted to JIMB; SNSF P400PB 186777, granted to XM).

## Data Availability Statemanet

The new DNA sequences used in this study are available from the NCBI (Supporting file 3, Table S3). Complete concatenated alignment (except new sequences generated in the present work) is available from Jardim de Queiroz and colleagues (Jardim de Queiroz et al., 2020) DOI: 10.17632/wccvm8p5gx.1. XML files to reconstruct and calibrate the phylogeny, and to reconstruct habitat preference are available from Supporting file 1. Custom codes used to reconstruct the biogeography with RevBayes are available from Supporting file 2 and from the GIT repository https://bitbucket.org/XavMeyer/biogeographyrevscript/src/master/.

## Author Contribution

LJQ and JIMB designed the research, analyzed and interpreted the results and drafted the manuscript. LJQ assembled the data and ran the analyses. IAB conducted part of the wet-laboratory routine. XM wrote the helper script specifying RevBayes analyses, helped to run the biogeographic analyses and to interpret the results. YC, RC, GTV, TEP and PAB provided samples and/or laboratorial infrastructure to generate part of the data. All authors read, contributed to the content and approved the final version.

## Supporting information (doi:10.17632/2fh5rj2zvg.1)

**Supporting file 1:** XML files for phylogenetic inference and tree calibration, and resulting trees in nexus format. These files correspond to four different runs (four scenarios as described in Table 1 in the main article). XML file for ancestral habitat reconstruction and resulting tree in nexus format are also available.

**Supporting file 2:** RevBayes scripts that were modified to constraint a given list of geographic ranges.

**Supporting file 3. Tables S1–S3. Table S1**. Marginal likelihoods obtained from the distinct analyses conducted with RevBayes to reconstruct ancestral ranges. **Table S2.**

Biohydrogeographic regions and their connectivity through time periods, according to our time-stratified model (M3). **Table S3.** Accession numbers of sequences generated in the present work.

